# Chimeric CRISPR guides enhance Cas9 target specificity

**DOI:** 10.1101/147686

**Authors:** Noah Jakimo, Pranam Chatterjee, Joseph M Jacobson

## Abstract

Oligonucleotide-guided nucleases (OGNs) have enabled rapid advances in precision genome engineering. Though much effort has gone into characterizing and mitigating mismatch tolerance for the most widely adopted OGN, *Streptococcus pyogenes* Cas9 (SpCas9), potential off-target interactions may still limit applications where on-target specificity is critical. Here we present a new axis to control mismatch sensitivity along the recognition-conferring spacer sequence of SpCas9’s guide RNA (gRNA). We introduce mismatch-evading lowered-thermostability guides (melt-guides) and exhibit how nucleotide-type substitutions in the spacer can reduce cleavage of sequences mismatched by as few as a single base pair. Cotransfecting melt-guides into human cell culture with an exonuclease involved in DNA repair, we observe indel formation on a standard genomic target at approximately 70% the rate of canonical gRNA and undetectable on off-target data.

---

The recent discoveries, characterizations, and modifications of natural oligonucleotide-guided nucleases associated with CRISPR and RNAi have empowered a genome-editing revolution ^1–4^. Low barriers for OGNs’ cost and design drive their widespread adoption over alternatives, including modular base-recognition domains (i.e., transcription activator like effector, zinc finger, and pumilio assemblies), which can be hard to synthesize, or meganucleases, which are difficult to engineer for new targets ^5–8^. Unlike protein-directed systems, OGNs also permit employing predictable nucleic acid chemistry and biophysics to alter native features ^9–12^.

Among the most important properties dictating the usage of a nucleic acid recognition system is its specificity. Thus, the desire to identify new methods diminishing potentially toxic or detrimental off-target activity has prompted many to measure and improve mismatch discrimination for RNA-guided SpCas9 - the most prevalent OGN and henceforth referred to as Cas9 ^13–15^. Up to now, others have increased its precision through broad approaches, such as controlling duration of exposure, enforcing co-localization on adjacent targets, or destabilizing binding affinity by minor variation ^16–19^. Here we present chimeric mismatch-evading lowered-thermostability guides that replace most gRNA spacer positions with DNA bases to suppress mismatched targets under Cas9’s catalytic threshold.

In this work, we confirm by *in vitro* cleavage assays that melt-guides can direct Cas9 with substantially enhanced mismatch discrimination. Moreover, we verify *in vivo* that melt-guides can achieve efficient mutagenesis with greater precision by providing deep sequencing data from transfected HEK293T cells stably expressing Cas9.

## DNA substitutions in Cas9 gRNA improve mismatch sensitivity

Efforts that have measured and modeled Cas9 target recognition imply a mechanism that includes incremental strand invasion between gRNA spacer and target sequence ^20, 21^. After prerequisite binding to a short protospacer adjacent motif (PAM), Cas9 helps stabilize DNA unwinding at a potential target as guide displaces its DNA:DNA base-pairs with RNA:DNA base-pairs (**Figure 1A**) ^22^. After the resulting structure, called an R-loop, expands beyond a ^~^15 base-pair exchange, Cas9 can then create a double-strand DNA break ^23, 24^.

**Figure 1.**
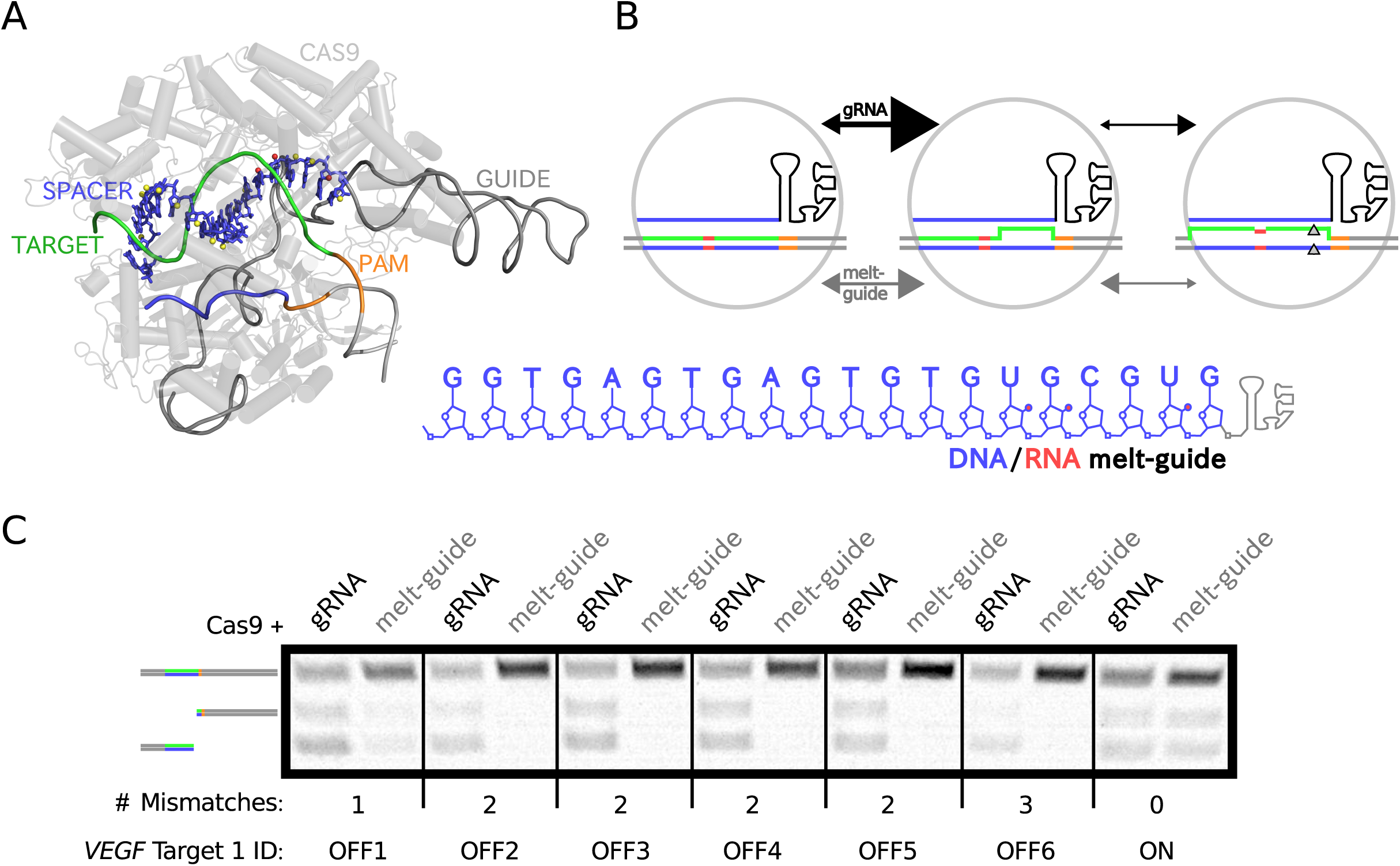
A Cas9 guide with DNA substitutions has reduced activity on mismatched targets. (a) Annotated 3D structure of a target-guide-Cas9 R-loop based on PDB 5F9R shown above the 2D structure of a melt-guide. Red and yellow spheres highlight RNA hydroxls eliminated and retained, respectively, in initial melt-guide designs. (b) Proposed model of relative R-loop expansion rate differences (represented by arrow sizes and directions) that increase mismatch sensitivity for melt-guides compared to gRNA. Red segments indicate mismatches between guide and target. (c) Inverted contrast-adjusted gel image of 4-hour Cas9 *in vitro* digests of targets with mismatches ranging from 0 to 3 using gRNA or melt-guide.

Motivated by studies on RNA/DNA chimera hybridization indicating more DNA content generally decreased duplex stability, we rationally designed chimeric melt-guides promoting the rehybridization of mismatched R-loops (**Figure 1B**) ^25, 26^. As illustrated, we selected candidate DNA-tolerant positions in gRNA by excluding most positions containing RNA-specific 2’-hydroxyl contacts with Cas9 that may help maintain assembly of active OGN. We confirmed *in silico* via Rosetta that our selection strategy had a proportionally greater energy score penalty on published target-bound structures than for unbound guide-Cas9 (**Figure 1 Supplement 1**) ^27^. Our interpretation that these scores, together, approximate R-loop stability and Cas9-guide affinity, emboldened us to substitute most gRNA spacer bases with DNA.

For a standard target sequence from human *VEGF*, we used commercially synthesized chimeric melt-guides and corresponding on- and off-target DNA substrates to compare a melt-guide’s mismatch discrimination to canonical gRNA when directing DNA cleavage. **Figure 1C** shows that a melt-guide containing 17 DNA bases was functional in a 4-hour digestion assay with purified Cas9 and produced 74% the amount of cleaved on-target substrate as did gRNA. The same melt-guide resulted in no detectable cleavage for all surveyed two-mismatch off-targets, which, in many cases gRNA-Cas9 cut faster than on-target substrate. Furthermore, on a challenging single-mismatch substrate that has been reported to be just as frequently an off-target for wild-type and high-fidelity Cas9 (hfCas9) variants, melt-guide reduced the digested fraction by four-fold ^18, 28^.

Additional *in vitro* assays demonstrate the generality of designing melt-guides for different genomic targets, but likewise reveal that targets comprising high GC and/or pyrimidine target content can limit sufficient destabilization to avoid cutting certain multi-mismatched sequences, even with melt-guides containing only DNA in the spacer (**Figure 1 Supplement 2**) ^29^. This limitation can be used to inform target-selection for a given application or it can be potentially overcome through combination of orthogonal destabilization techniques, such as truncating guide or complexing it with hfCas9. We have identified other nucleotide-type substitutions that also enhance specificity, including unlinked nucleic acid (UNA) and abasic or universal base nucleotides at sequence positions with low-priority or no mismatches in the ensemble of possible off-targets (**Figure 1 Supplement 3**) ^30, 31^.

## R-loop expansion kinetics determine melt-guide specificity

Whereas Cas9 is known to rapidly cleave DNA, its rates with melt-guides slowed appreciably (**Figure 2A**) ^32^. In order to confirm R-loop expansion contributes more than mismatched hybridization to this change in kinetics, we ran time-coursed digestions using substrates that were either double-stranded (ds) or single-stranded (ss) along the target (**Figure 2B**). Within minutes, Cas9 with canonical gRNA was able to cut both ds- and ss- target to near completion. For melt-guide-directed cleavage, we instead observed steady digestion of no-mismatch ds-targets over several hours, yet rates on ss-targets about as rapid as gRNA’s and at similar timescales in the presence and absence of mismatches. The fast error-prone cuts we detected upon removing strand-displacement from cleavage dynamics support that R-loop destabilization contributes mainly to melt-guides’ improved specificity.

**Figure 2.**
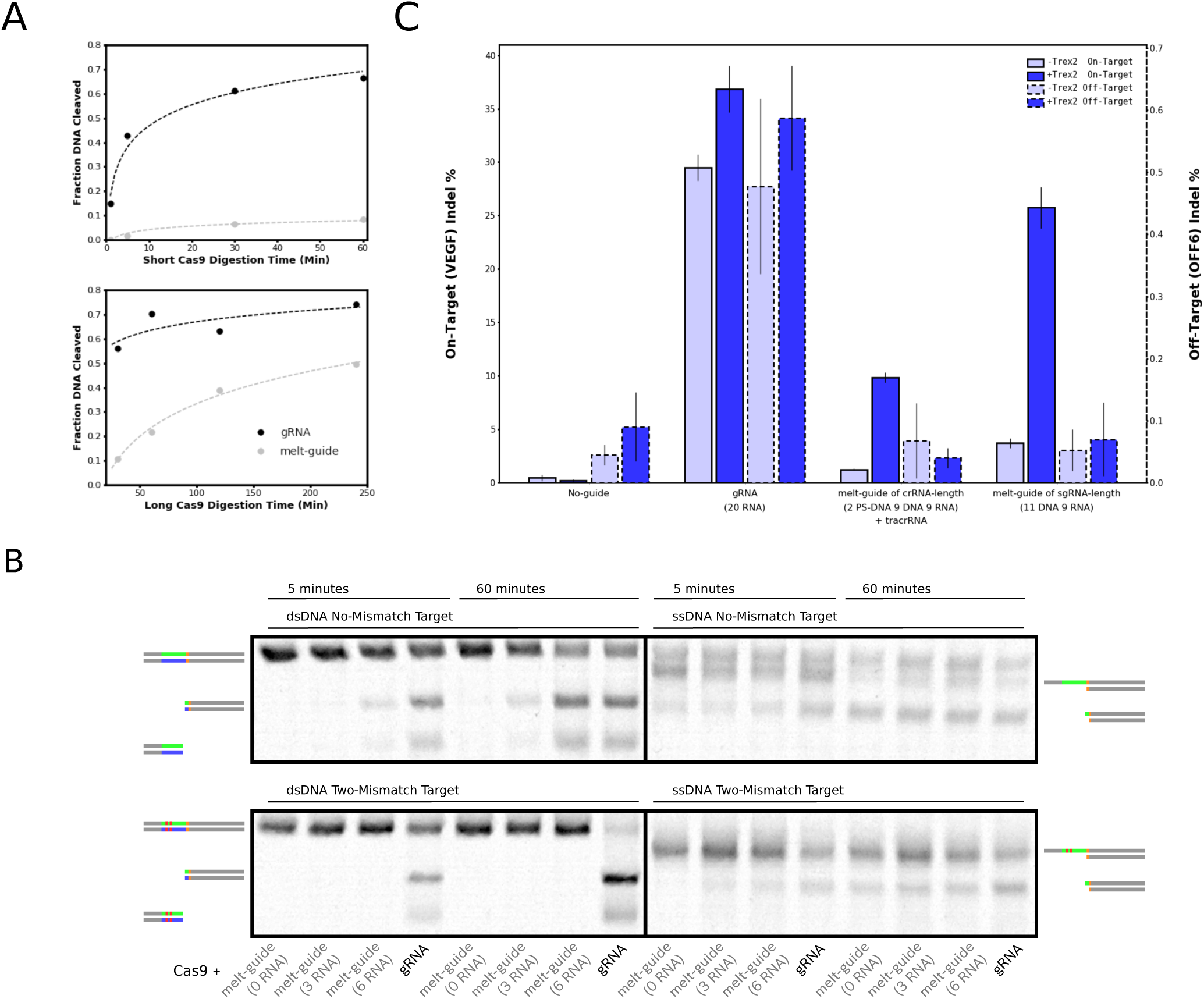
Melt-guide strand invasion determines DNA cleavage and gene-editing rates. (a) Melt-guide (gray) and gRNA (black) cleavage time courses of no-mismatch target with Cas9 plot alongside least-squares-fitting logarithmic functions (dashed curves). (b) Inverted contrast-adjusted gel images of short (left-side within each quadrant) and long (right-side within each quadrant) Cas9 digests of double-stranded (left-half) and single-stranded (right-half) targets with two mismatches (bottom) or none (top) using gRNA or melt-guides with spacers containing either all-DNA, 3 DNA distributed as previously described, or an additional 3 DNA fill-in. (c) Dual y-axis chart shows deep sequencing indel measurements on-target and at a known off-target, comparing mutagenesis by gRNA to melt-guides designed with DNA substitutions in their first 11 positions with and without Trex2 overexpression (blue and light blue, respectively). Nucleic acid-type content in the guide’s spacer is noted in parentheses.

Future single-molecule fluorescent resonance energy transfer (FRET) measurements of melt-guides can be used to obtain finer detail of recognition kinetics and Cas9 conformational changes, complementary to previous work using gRNA ^33, 34^. While we noticed melt-guides that include all-DNA-spacer did not introduce drastic structural changes that would have prevented cleavage, it is unclear whether such guides more closely adopt A-form or B-form duplexes with their target. This uncertainty arises from antagonistic influences of Cas9 pre-loading guide in an unpaired A-form versus the favored B-forming tendency of DNA:DNA dimers ^27, 35^. The exact extent to which the helicity is altered for melt-guides in oligonucleotide-protein complexes could be solved from a crystal structure of the bound melt-guide OGN.

## Melt-guide and Trex2 co-transfection reduces off-target genome editing

To test the use of melt-guides for genome editing, we transfected *VEGF*-targeting melt-guide oligos into HEK293T cells stably expressing Cas9 and enzymatically measured insertion/deletion (indel) mutations (**Figure 2 Supplement 1**). Initial attempts yielded unsatisfactorily low mutagenesis, which we believe resulted from unfavorable relative rates of: (i) guide oligo degradation, (ii) slower R-loop expansion, and (iii) errorless non-homologous end-joining (NHEJ) repair ^36, 37^. We tried counteracting degradation with oligo lifetime-lengthening modications (e.g., phosphorothioate (PS-DNA) or inverted terminal bases and 2’-O-methyl RNA substitutions on non-spacer guide positions) and partially restored cleavage rates by using fewer DNA substitutions in melt-guides ^9^. Since these tactics did not lead to substantial improvement, we later pursued methods that could bias genomic double-strand breaks towards more error-prone repair.

Overexpression of the mammalian 3’ exonuclease Trex2, associated with DNA damage processing, has been reported to raise indel rates for various sequence-specific gene editing systems without causing toxicity ^38–40^. Therefore, we added Trex2 expression plasmid to transfections and measured effected mutations by deep sequencing (**Figure 2C**) ^41^. We found a melt-guide containing mostly DNA in spacer bases produced indel percentages above 25% on-target, which acceptably translates to ~70% gRNA’s rate. Crucially, on an off-target where we detected gRNA-induced mutations, melt-guides’ indel percentages fell below our no-guide negative control. Between melt-guide types, single-molecule gRNA (sgRNA) length melt-guides consistently generated more than double the indel rate of melt-guides derived from shorter CRISPR RNA (crRNA) sequence, which need to duplex with trans-activating crRNA (tracrRNA). Despite Trex2 addition increasing indel percentages roughly seven-fold for both melt-guide types, the exonuclease had marginal impact on gRNA-directed mutation rates.

Others have achieved enhanced Cas9 specificity and could maintain high indel rates on-target without an accessory exonuclease ^18, 28^. However, their experiments relied on transcribing all OGN components to abundant cellular concentrations. On one hand, a similar Trex2 supplementation strategy may benefit applications where some components are delivered as oligo or protein - which may include DNA-guided editing with Argonaute ^42, 43^. On the other hand, a reverse-transcribable melt-guide with only DNA bases could lessen dependence on Trex2 for efficient mutagenesis. Towards that end, we show *in vitro* cleavage directed by tracrRNA in duplex with a crRNA-length melt-guide containing a single RNA outside of the spacer sequence (**Figure 2 Supplement 2**). Chimeras with such sparse RNA content are furthermore likely resistant to most RNases.

## Conclusion

In the case of Cas9, we improve the precision of target activity *in vitro* and *in vivo* with mismatch-evading lowered-thermostability guides. We believe melt-guides should be extensible to the expanding collection of CRISPR systems by extrapolating either from chimeric oligo libraries to scan nucleotide-type substitution or from published crystal structure data to avoid disrupting RNA-specific interactions (i.e., Cpf1 guide’s pseudoknots) ^44, 45^. Given the minimal RNA content that we found to be sufficient for guiding Cas9, additional protein engineering - perhaps through homolog alignments - may enable the realization of all-DNA melt-guides.

## Methods

### Cas9-guide *in vitro* DNA digestions

Mixed nucleotide-type and RNA oligos, designed as Cas9 guides for selected standard genomic targets, were obtained from Integrated DNA Technologies (IDT). A 1 M dilution was prepared for stocks of guide derived from sgRNA or crRNA and the latter was combined with equimolar tracrRNA (GE Dharmacon). Reactions consisted of 20 M pre-annealed guide stock, 20 nM purified Cas9 from New England BioLabs (NEB), 10x NEB reaction buffer, and 500 g of IDT-synthesized dsDNA target in 30 1 mixes. Samples were incubated at 37C and digested products separated by TAE-gel electrophoresis. Images of cleaved fractions from SYBR-Safe dsDNA gel stain (Thermo Fisher) under a blue light lamp were quantified using ImageJ software.

### Preparation of single-stranded target DNA substrates

Target substrates were PCR-amplified using a primer oligo set (IDT) with 5’ phosphorylation for only the primer generating PAM-sided strands. Amplicons purified on anion-resin exchange columns (Qiagen) were digested by Lambda exonuclease (NEB), a 5’-to-3’ enzyme that prefers phophorylated ends of dsDNA, to yield ssDNA of the strand opposite of PAM. Following subsequent column purification, ssDNAs were annealed to a primer beginning at the PAM site of the removed strand and templated for extension by DNA polymerase (NEB).

### Genomic indel production and measurements

HEK293T cells stably expressing Cas9 purchased from GeneCopoeia were plated to 250,000 cells / 35 mm well in 2.5 ml Dulbeccos Modified Eagles Medium with 10% Fetal Bovine Serum and incubated at 37 C and 5% CO_2_. The next day, transfections via TransIT-X2 reagent (Mirus Bio) delivered a 25 nM final concentration of guide with or without 2.5 g pExodus CMV.Trex2, which was a gift from Dr. Andrew Scharenberg (Addgene plasmid #40210). After an additional 48 hours, genomic DNA was isolated using Epicentre QuickExtract solution and indel production was visualized by a common T7 Endonuclease I assay (NEB) on amplicons from on-target and known off-target regions ^46^. Amplicons were then prepared for deep sequencing with Nextera-XT tagmentation (Illumina) and run on a MiSeq 2x300 v3 kit (Illumina). Reads were analyzed using the CRISPResso software pipeline for precise indel percentages from biological and technical duplicates ^41^.

**Table 1:**
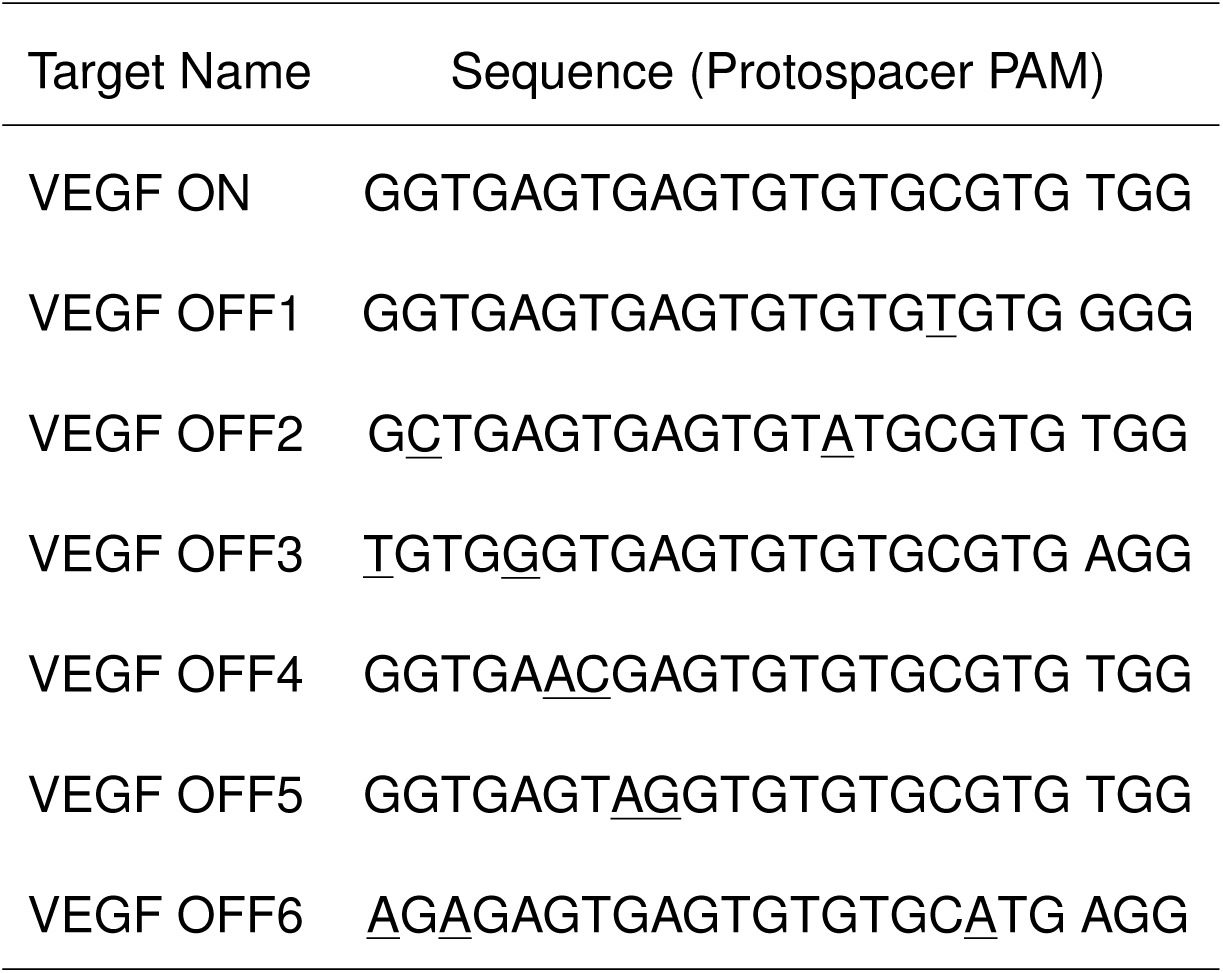
Sequence information with underlined mismatches.

## Acknowledgements

We thank Thrasyvolous Karydis and Frank Soucy for computational analysis of Cas9-guide structures and Keren Greenbaum for technical support. We thank Stuart Levine and the MIT BioMicroCenter for Illumina library prep and sequencing. We also thank Dr. Ed Boyden for access to cell culture, in addition to Dr. Neil Gershenfeld and Dr. Shuguang Zhang for shared lab equipment.

## Author Contributions

NJ, Conceptualization, *In vitro* and *In vivo* Investigation, Data Analysis, Writingoriginal draft, Writingreview and editing; PC, *In vivo* Investigation, Writingreview; JMJ, Conceptualization, Writingreview and editing

---

**Supplementary Figure 1.1.** Rosetta energy scores with DNA substitutions in bound and unbound structures from PDBs 4UN3 and 4ZT0.

**Supplementary Figure 1.1.** Gel images from Cas9 digests with melt-guides of on and off-target sequences for *EMX*, *FANCF*, and *VEGF2*.

**Supplementary Figure 1.3.** Gel images from Cas9 digests with UNA substitutions in a melt-guide that targets *VEGF2*.

**Supplementary Figure 2.1.** T7EI endonuclease assay on genomic *VEGF* amplicons upon Cas9 mutagenesis with various melt-guide designs.

**Supplementary Figure 2.2.** Inverted gel image of Cas9 digest of no-mismatch *VEGF* target with almost all-DNA melt-guide of crRNA-length.

## References

1. Jinek, M. et al. Rna-programmed genome editing in human cells. eLife 2, e00471 (2013). Original DateCompleted: 20130207, Original DateCompleted: 20140206.

2. Mali, P. et al. Rna-guided human genome engineering via cas9. Science (New York, N.Y.) 339, 823–826 (2013).

3. Cong, L. et al. Multiplex genome engineering using crispr/cas systems. Science (New York, N.Y.) 339, 819–823 (2013).

4. Komor, A. C., Badran, A. H. & Liu, D. R. Crispr-based technologies for the manipulation of eukaryotic genomes. Cell 168, 20–36 (2017).

5. Reyon, D. et al. Flash assembly of talens for high-throughput genome editing. Nature biotechnology 30, 460–465 (2012).

6. Ramirez, C. L. et al. Unexpected failure rates for modular assembly of engineered zinc fingers. Nature methods 5, 374–375 (2008).

7. Adamala, K. P., Martin-Alarcon, D. A. & Boyden, E. S. Programmable rna-binding protein composed of repeats of a single modular unit. Proceedings of the National Academy of Sciences 113, E2579–E2588 (2016).

8. Takeuchi, R., Choi, M. & Stoddard, B. L. Redesign of extensive protein–dna interfaces of meganucleases using iterative cycles of in vitro compartmentalization. Proceedings of the National Academy of Sciences 111, 4061–4066 (2014).

9. Hendel, A. et al. Chemically modified guide rnas enhance crispr-cas genome editing in human primary cells. Nature biotechnology 33, 985–989 (2015).

10. Jain, P. K. et al. Development of light-activated crispr using guide rnas with photocleavable protectors. Angewandte Chemie (International ed. in English) 55, 12440–12444 (2016).

11. Lee, K. et al. Synthetically modified guide rna and donor dna are a versatile platform for crispr-cas9 engineering. eLife 6 (2017).

12. Ui-Tei, K. et al. Functional dissection of sirna sequence by systematic dna substitution: modified sirna with a dna seed arm is a powerful tool for mammalian gene silencing with significantly reduced off-target effect. Nucleic acids research 36, 2136–2151 (2008).

13. Schaefer, K. A. et al. Unexpected mutations after crispr-cas9 editing in vivo. Nature methods 14, 547–548 (2017).

14. Tsai, S. Q. et al. Circle-seq: a highly sensitive in vitro screen for genome-wide crispr-cas9 nuclease off-targets. Nature methods 14, 607–614 (2017).

15. Doench, J. G. et al. Optimized sgrna design to maximize activity and minimize off-target effects of crispr-cas9. Nature biotechnology 34, 184–191 (2016).

16. Davis, K. M., Pattanayak, V., Thompson, D. B., Zuris, J. A. & Liu, D. R. Small molecule-triggered cas9 protein with improved genome-editing specificity. Nature chemical biology 11, 316–318 (2015).

17. Ran, F. A. et al. Double nicking by rna-guided crispr cas9 for enhanced genome editing specificity. Cell 154, 1380–1389 (2013).

18. Kleinstiver, B. P. et al. High-fidelity crispr-cas9 nucleases with no detectable genome-wide off-target effects. Nature 529, 490–495 (2016).

19. Fu, Y., Reyon, D. & Joung, J. K. Targeted genome editing in human cells using crispr/cas nucleases and truncated guide rnas. Methods in enzymology 546, 21–45 (2014).

20. Farasat, I. & Salis, H. M. A biophysical model of crispr/cas9 activity for rational design of genome editing and gene regulation. PLoS computational biology 12, e1004724 (2016).

21. Josephs, E. A. et al. Structure and specificity of the RNA-guided endonuclease cas9 during DNA interrogation, target binding and cleavage. Nucleic Acids Research 43, 8924–8941 (2015).

22. Jiang, F., Zhou, K., Ma, L., Gressel, S. & Doudna, J. A. Structural biology. a cas9-guide rna complex preorganized for target dna recognition. Science (New York, N.Y.) 348, 1477–1481 (2015).

23. Jiang, F. et al. Structures of a crispr-cas9 r-loop complex primed for dna cleavage. Science (New York, N.Y.) 351, 867–871 (2016).

24. Kiani, S. et al. Cas9 grna engineering for genome editing, activation and repression. Nature methods 12, 1051–1054 (2015).

25. Sugimoto, N. et al. Thermodynamic parameters to predict stability of rna/dna hybrid duplexes. Biochemistry 34, 11211–11216 (1995).

26. Nakano, S.-i., Kanzaki, T. & Sugimoto, N. Influences of ribonucleotide on a duplex conformation and its thermal stability: study with the chimeric rna-dna strands. Journal of the American Chemical Society 126, 1088–1095 (2004).

27. Nishimasu, H. et al. Crystal structure of cas9 in complex with guide rna and target dna. Cell 156, 935–949 (2014).

28. Slaymaker, I. M. et al. Rationally engineered cas9 nucleases with improved specificity. Science (New York, N.Y.) 351, 84–88 (2016).

29. Gyi, J. I., Conn, G. L., Lane, A. N. & Brown, T. Comparison of the thermodynamic stabilities and solution conformations of dna.rna hybrids containing purine-rich and pyrimidine-rich strands with dna and rna duplexes. Biochemistry 35, 12538–12548 (1996).

30. Snead, N. M., Escamilla-Powers, J. R., Rossi, J. J. & McCaffrey, A. P. 5’ unlocked nucleic acid modification improves sirna targeting. Molecular therapy. Nucleic acids 2, e103 (2013).

31. Zhang, J. et al. Modification of the sirna passenger strand by 5-nitroindole dramatically reduces its off-target effects. Chembiochem: a European journal of chemical biology 13, 1940–1945 (2012).

32. Sternberg, S. H., LaFrance, B., Kaplan, M. & Doudna, J. A. Conformational control of dna target cleavage by crispr-cas9. Nature 527, 110–113 (2015).

33. Singh, D., Sternberg, S. H., Fei, J., Doudna, J. A. & Ha, T. Real-time observation of dna recognition and rejection by the rna-guided endonuclease cas9. Nature communications 7, 12778 (2016).

34. Szczelkun, M. D. et al. Direct observation of r-loop formation by single rna-guided cas9 and cascade effector complexes. Proceedings of the National Academy of Sciences of the United States of America 111, 9798–9803 (2014).

35. Gyi, J. I., Lane, A. N., Conn, G. L. & Brown, T. Solution structures of dna.rna hybrids with purine-rich and pyrimidine-rich strands: comparison with the homologous dna and rna duplexes. Biochemistry 37, 73–80 (1998).

36. Suzuki, K. et al. In vivo genome editing via crispr/cas9 mediated homology-independent targeted integration. Nature 540, 144149 (2016). URL http://dx.doi.org/10.1038/nature20565.

37. Schmid-Burgk, J. L., Hning, K., Ebert, T. S. & Hornung, V. Crispaint allows modular base-specific gene tagging using a ligase-4-dependent mechanism. Nature communications 7, 12338 (2016).

38. Delacte, F. et al. High frequency targeted mutagenesis using engineered endonucleases and dna-end processing enzymes. PloS one 8, e53217 (2013).

39. Certo, M. T. et al. Coupling endonucleases with dna end-processing enzymes to drive gene disruption. Nature methods 9, 973–975 (2012).

40. Chari, R., Mali, P., Moosburner, M. & Church, G. M. Unraveling crispr-cas9 genome engineering parameters via a library-on-library approach. Nature methods 12, 823–826 (2015).

41. Pinello, L. et al. Analyzing crispr genome-editing experiments with crispresso. Nature biotechnology 34, 695–697 (2016).

42. Enghiad, B. & Zhao, H. Programmable dna-guided artificial restriction enzymes. ACS synthetic biology 6, 752–757 (2017).

43. Lee, S. H. et al. Failure to detect dna-guided genome editing using natronobacterium gregoryi argonaute. Nature biotechnology 35, 17–18 (2016).

44. Yamano, T. et al. Crystal structure of cpf1 in complex with guide rna and target dna. Cell 165, 949–962 (2016).

45. Burstein, D. et al. New crispr-cas systems from uncultivated microbes. Nature 542, 237–241 (2017).

46. Vouillot, L., Thlie, A. & Pollet, N. Comparison of t7e1 and surveyor mismatch cleavage assays to detect mutations triggered by engineered nucleases. G3 (Bethesda, Md.) 5, 407–415 (2015).

